# Phylogenomic Insights into the Origin of Primary Plastids

**DOI:** 10.1101/2020.08.03.231043

**Authors:** Iker Irisarri, Jürgen F. H. Strassert, Fabien Burki

## Abstract

The origin of plastids was a major evolutionary event that paved the way for an astonishing diversification of photosynthetic eukaryotes. Plastids originated by endosymbiosis between a heterotrophic eukaryotic host and a cyanobacterium, presumably in a common ancestor of all primary photosynthetic eukaryotes (Archaeplastida). A single origin of primary plastids is well supported by plastid evidence but not by nuclear phylogenomic analyses, which have consistently failed to recover the monophyly of Archaeplastida hosts. Importantly, the monophyly of both plastid and host (nuclear) genomes is required to support a single ancestral endosymbiosis, whereas non-monophyletic hosts could be explained under scenarios of independent or serial eukaryote-to-eukaryote endosymbioses. Here, we assessed the strength of the signal for the Archaeplastida host monophyly in four available phylogenomic datasets. The effect of phylogenetic methodology, data quality, alignment trimming strategy, gene and taxon sampling, and the presence of outlier genes loci were investigated. Our analyses revealed a general lack of support for host monophyly in the shorter individual datasets. However, when analyzed together under rigorous data curation and complex mixture evolutionary models, the combined dataset consistently recovered the monophyly of Archaeplastida hosts. This study represents an important step towards better understanding the eukaryotic evolution and the origin of plastids.

## Introduction

The origin of plastids by endosymbiosis with cyanobacteria was a key evolutionary event that allowed eukaryotes to perform oxygenic photosynthesis (i.e., photoautotrophy). This innovation paved the way for an astonishing diversification of micro- and macroscopic algae and land plants in all sunlit environments. Generally, plastids have been viewed to originate from a single endosymbiotic event in a common ancestor of three well-defined lineages: Glaucophyta, Rhodophyta (red algae), and Chloroplastida (green algae and land plants), collectively known as Archaeplastida (Adl et al. 2005) or Plantae (Cavalier-Smith 1998). These lineages harbor primary plastids, i.e. derived by endosymbiosis directly from cyanobacteria, and thereafter will be referred to as primary photosynthetic eukaryotes (PPE). Despite structural and genomic differences, the plastids of Glaucophyta, Rhodophyta, and Chloroplastida share many similarities such as homologous protein transport apparatus, gene content, and synteny blocks that have been interpreted as evidence supporting their common origin (Cavalier-Smith 2000; Löffelhardt 2014; Mackiewicz and Gagat 2014). Furthermore, these observations are consistent with the monophyly of PPE generally recovered in plastid phylogenies.

Unlike plastid phylogenies, resolving the relationship among host (nuclear) lineages has proven more difficult and the question of the monophyly of Archaeplastida currently stands out as one of the major knowledge gaps in the eukaryotic Tree of Life (Burki et al. 2020). This is because inferring the relationships among PPE necessarily involves resolving the relationships among other deep-branching eukaryotic groups such as Cryptophyta or Haptophyta, all notoriously difficult to place in the eukaryotic tree (Burki et al. 2020). With a few exceptions (Lax et al. 2018; Price et al. 2019), the majority of nuclear phylogenomic studies have not recovered the monophyly of PPE (Hampl et al. 2009; Baurain et al. 2010; Parfrey et al. 2010; Burki et al. 2012, 2016; Brown et al. 2013; Yabuki et al. 2014; Janouškovec et al. 2017), or did so with low statistical support (Burki et al. 2007, 2009), or using a very sparse taxon sampling (Rodríguez-Ezpeleta et al. 2005, 2007a; Deschamps and Moreira 2009). Recent studies reported contradicting Bayesian and maximum likelihood trees regarding Archaeplastida monophyly (Brown et al. 2018; Gawryluk et al. 2019; Strassert et al. 2019). Importantly, the majority of studies that did not to recover monophyletic PPE did not converge to a robust alternative topology either, leading to the current situation where the monophyly of Archaeplastida is clear from plastids but inconclusive from host data.

While cell biological and genomic characters in primary plastids are more easily explained under the hypothesis of a single endosymbiosis, the observed similarities could at face value be the result of parallel or convergent evolution (Stiller 2014). In fact, the monophyly of plastids is a necessary but insufficient condition to invoke a single endosymbiosis in the ancestor of Archaeplastida. Alternative biological scenarios can explain plastid monophyly when hosts are not monophyletic, including (i) serial endosymbioses or (ii) the independent acquisition of plastids from closely-related (and now extinct) cyanobacterial lineages (Supplementary Fig. 1; Mackiewicz and Gagat 2014). Therefore, the common origin of primary plastids in the ancestor of PPE can only be hypothesized if both plastid and host lineages are shown to be monophyletic.

Here, we assessed the evidence for the monophyly of Archaeplastida by investigating the signal and conflict in four available nuclear phylogenomic datasets. After correcting for systematic biases, none of the datasets supported the monophyly of Archaeplastida, showing only diverse and weakly supported relationships. In search for the reasons of this lack of signal, we investigated the effects of phylogenetic inference methods and models, gene sampling, taxon sampling, and the presence of outlier loci. To overcome the limitations observed in the four source datasets, we generated six combined datasets after rigorous data curation and investigated the effect of data quality, systematic errors (model misspecifications), and the effect of alignment trimming algorithms in recovering deep eukaryotic relationships. When analyzed under complex mixture models, the combined datasets provided a congruent hypothesis for deep eukaryotic relationships with monophyletic Archaeplastida.

## Materials and Methods

### Published Phylogenomic Datasets

We chose four published representative datasets that were assembled independently: (Baurain et al. 2010; BAU), (Brown et al. 2013; BRO), (Katz and Grant 2015; KAT), (Burki et al. 2016; BUR). All datasets consisted on concatenated protein alignments, except for KAT that additionally contained the 18S rRNA gene. In KAT, protein and DNA alignments were analyzed both jointly and independently. All datasets were taken “as-is” from the original authors with minimal intervention to standardize taxonomic names. Because the concatenated alignment in KAT did not retain gene boundary information, we re-created the protein dataset following their published protocol: the eukaryotic-only gene alignments provided by the authors (https://datadryad.org/resource/doi:10.5061/dryad.db78g.2/13.2) were subsampled for the 231 eukaryotic taxa and 149 protein alignments used in their final tree (Fig. 1 of Katz and Grant [2015]) and alignment columns with >50% missing data were removed, followed by concatenation. The completeness of all four datasets was assessed with AliStat v.1.11 (Wong et al. 2014) using four recently proposed metrics (Wong et al. 2020).

### Maximum Likelihood Re-analysis

To account for the effect of inference methods and models used in the original studies, the four datasets were analyzed under equivalent conditions: maximum likelihood (ML) under best-fit site-homogeneous (LG+F+Γ4) and site-heterogeneous (LG+C40/C60+F+Γ4) models using IQTREE v.1.5.4 (Nguyen et al. 2015). Best-fit models were selected with ModelFinder (Kalyaanamoorthy et al. 2017) and branch support was assessed with 1,000 pseudo-replicates of ultrafast bootstrapping (UFBoot; Hoang et al. 2017). The two shorter datasets (BAU, BRO) used the C40 empirical mixture to avoid overparameterization (increasing the number of profiles was not supported because some profiles had zero weights).

### Comparison of Gene and Taxon Sampling Across Datasets

The effect of gene and taxon sampling was assessed by comparing ML trees from data subsets of shared genes and taxa across datasets. We considered all six possible pairwise gene and taxon overlaps between datasets plus a seventh one as the intersection of all four. To identify shared genes, we standardized gene names to that of the human ortholog, which was identified by BLASTP against all human proteins (GRCh38.p7; ENSEMBL 87) using human (or metazoan) sequences in gene alignment as queries (or the longest sequence if the former covered <50% of the protein). To identify shared taxa, taxon names were also standardized. A total of 16 “gene overlap” datasets were constructed as follows: for each set of pairwise shared genes, two gene overlap datasets were assembled by subsampling the two source datasets (i.e., same gene name, different source alignment); for the set of genes shared across all four datasets, four such datasets were built. The analogous procedure was used to create 16 “taxon overlap” datasets. Therefore, “gene overlap” datasets contained equivalent genes but retained the original taxon sampling, whereas “taxon overlap” datasets had comparable taxon sampling but retained the original gene sampling. Because BAU and BUR datasets contained some chimeric taxa from closely-related species (10 and 14, respectively), taxon sets were considered to overlap whenever at least one of the species in the chimeric taxa matched.

The 32 datasets were analyzed by ML using IQ-TREE v.1.5.1 under best-fit site-homogeneous models and 1,000 UFBoot pseudo-replicates. The eight datasets representing subsets of genes and taxa shared by all four datasets were further analyzed under the site-heterogeneous LG+C60+F+Γ4 model. The topological distance among all resulting ML trees were measured with normalized Robinson-Foulds distances (nRF = RF/RFmax; Kupczok et al. 2008), which accounts for differences in taxon sampling, using ETE 3 (Huerta-Cepas et al. 2016). Pairwise nRF distances were visualized using Kruskal’s non-metric multidimensional scaling in the R package MASS (Venables and Ripley 2002).

### Quantifying Gene Support for Archaeplastida Monophyly

We assessed the support for PPE monophyly in the four original datasets following Shen et al. (2017). Briefly, we calculated the gene-wise log-likelihood score differences (ΔGLS) between tree topologies that differ in the monophyly of PPE or lack thereof.

For BAU, BRO, and BUR, the unconstrained ML trees inferred above had non-monophyletic PPE and the alternative trees were built by constrained ML searches (IQTREE; LG+C40/60+F+Γ4). For KAT, the unconstrained ML tree represented PPE monophyly and the alternative was built by breaking PPE monophyly according to BRO’s unconstrained tree (it recovered a higher likelihood than using BAU or BUR). Site-wise log-likelihoods were calculated in IQ-TREE under the above models; likelihood differences between the competing topologies were calculated per site and averaged per gene (ΔGLS) to quantify the gene support for or against Archaeplastida monophyly. For each dataset, we surveyed 5% of the most deviant genes (largest ΔGLS, for or against Archaeplastida) and manually inspected alignments and ML gene trees in search of obvious contaminations or paralogy problems. To confirm paralogy issues, nuclear, mitochondrial, and/or plastid homologs from reference eukaryotes (human, mouse, rice, *Arabidopsis*, yeast) were retrieved from UNIPROT and added into gene alignments. To assess the effect of identified paralogs in the overall tree topologies, we removed problematic sequences and re-inferred ML trees from entire datasets as specified above.

### New Combined Dataset

To test whether data combination can improve the resolution of deep eukaryotic relationships, we merged the four original datasets accounting for shared genes and taxa and reducing missing data using available genomes, transcriptomes, or ESTs from public databases: NCBI, ENSMBL, EukProt (Richter et al. 2020), MMETSP (Keeling et al. 2014), 1KP (Leebens-Mack et al. 2019), and OrthoMCL-DB v.5 (Chen et al. 2006). For most species, protein sets were publicly available. For five species, transcriptomes were assembled *de novo* using Trinity v.2.5.1 with default settings. For transcriptomes and ESTs, ORFs were predicted using TransDecoder v.3.0.1 (with the corresponding genetic code). The combined dataset was built in two steps. First, new taxa were added to the dataset containing the most genes (BUR, 250 genes). In practice, we started from a taxon-enriched version of the same dataset (Strassert et al. in prep) to which homologs for missing taxa were added using BLASTP (e-value <10^-6^) and retaining the two best hits to increase the chance of identifying the correct ortholog. The sets of homologous sequences were masked with PREQUAL v.1.01 (Whelan et al. 2018) (default settings, excluding fast-evolving taxa — prokaryotes, Ascetosporea, Hexamitidae, Microsporidia, and *Guillardia* nucleomorph — to avoid masking legitimate residues); aligned with MAFFT G-INS-i v.7.310 (Katoh and Standley 2013) with a variable scoring matrix to control for over-alignment (VSM; ‘--allowshift -- unalignlevel 0.6’; [Katoh and Standley 2016]); alignment columns with >99% missing data were trimmed with BMGE v.1.12 (Criscuolo and Gribaldo 2010); and gene trees were inferred with FastTree v.2.1.3 (Price et al. 2010) with default settings. Gene trees were visualized to ensure orthology, aided by the presence of appropriately labelled prokaryotic homologs and known paralogs from reference species. Obvious paralogs or contaminants were flagged, along with shorter homologs (up to two homologs per taxon were retained in BLASTP). All flagged sequences, as well as species not present in any of the four original datasets, were removed from the original (pre-PREQUAL) orthologous sets. The genes *FTSJ1* and *tubb* were excluded due to unresolved deep paralogy likely indicating a complex evolutionary history or too low phylogenetic signal.

In a second step, we targeted additional genes not present in BUR. Homologous gene alignments from BAU, BRO, and KAT were merged and complemented with sequences from missing taxa using BLASTP as above. To aid in the identification of contaminants and paralogs, several prokaryotic and eukaryotic homologs from reference genomes were added. Gene sets were masked and aligned as specified above, and filtered with Divvier (option ‘-partial’), which implements an HMM-based parametric model that allows removing residues from alignment columns that lack strong homology evidence (Ali et al. 2019). Gene trees were inferred using RAxML v.8.2.4 (Stamatakis 2014) under LG+Γ4 and 100 rapid bootstrap replicates; visualized for the identification of obvious paralogs and contaminants, which were removed along with duplicated taxa from the original (pre-PREQUAL) orthologus sets. A second round of masking, aligning, divvying, gene tree inference, and visualization was done in order to flag possible remaining contaminants or paralogs. During the two rounds of dataset cleanup a total of 28 genes were excluded due to unresolved deep paralogy, likely indicating complex evolutionary histories or low signal (including *EF2*, see below). The final gene sets were masked and aligned as specified above and concatenated with SCaFoS v.1.25 (Roure et al. 2007). The new combined dataset had 311 (248 + 63) genes and 344 taxa. Gene alignments of the combined dataset were subjected to three alignment trimming methods: (i) untrimmed (in practice, >99% incomplete columns removed), (ii) filtering non-homologous residues with Divvier (‘-partial’), and (iii) entropy-based block trimming with BMGE (‘-b 5 -m BLOSUM75 -g 0.2’). The sets of 311 gene alignments were concatenated into three datasets containing respectively (i) 202,042 (COMB-UNTRIM), (ii) 160,090 (COMB-DIVPART), and (i) 75,055 (COMB-BMGE) aligned amino acids, respectively. The combined (untrimmed) dataset was inspected for outlier loci by calculating gene-wise log-likelihoods and visualizing gene trees of the 5% most deviant genes, as done above.

ML trees were inferred from the three combined datasets using IQ-TREE v.1.6.5 with best-fit LG+I(+F)+Γ4 site-homogeneous models and 1,000 UFBoot pseudo-replicates. For computational tractability, we reduced the three datasets to 98 taxa (maintaining phylogenetic diversity) and analyzed them under more complex LG+C60+Γ4 mixture models. In addition, the shortest taxon-reduced dataset (COMB-BMGE-98) was analyzed under the better-fitting CAT-GTR model in PhyloBayes MPI v.1.8 (Lartillot et al. 2013) (larger datasets were intractable). The relative fit of LC60 vs. CAT-GTR was assessed by 10-fold cross-validation using PhyloBayes MPI and a random gene sample of 20,000 amino acid positions. The topological distance among the resulting ML and Bayesian trees, along with those of the original datasets, were calculated as nRF and plotted after non-metric multidimensional scaling.

### Effect of Alignment Filtering Algorithms

We approximated the phylogenetic signal in the three differently trimmed combined datasets by calculating the topological congruence between gene trees and the corresponding concatenated ML trees. Gene trees were inferred by ML in IQ-TREE v.1.6.10 with best-fit models and SH-aLRT with 1,000 replicates. For each treatment (untrimmed, Divvier partial, BMGE), we quantified the topological distances between gene trees and concatenated trees with nRF and the proportion of highly (SH-aLRT > 0.85) and lowly supported bipartitions that were congruent or not with the concatenated trees. Calculations were implemented in python3 with the aid of ETE3.

## Results

### Controlling for Evolutionary Model and Gene and Taxon Sampling Does Not Resolve Incongruence

A direct comparison of the phylogenetic relationships reported in the four original studies is complicated by the fact that they were analyzed with different methods and models, and only partially overlapped in genes and taxa. These differences make it difficult to identify the contribution of phylogenetic methodologies and of specific loci and taxa combinations in resolving the placement of PPE in the tree. To obtain baseline information across the four datasets, we re-analyzed the original alignments under consistent methods and models (IQ-TREE ML under best-fit site-homogeneous and mixture models). These analyses confirmed the recovery of monophyletic Archaeplastida in KAT (UFBoot = 91-95%) and lack thereof in the remaining three datasets, which did not converge to a congruent alternative topology (Fig. 1). The use of empirical profile mixture models (+C40/C60) did not change the inferred relationships among PPE.

Subsets of shared genes and taxa were equally inconclusive regarding the relationships among PPE. The monophylies of the three main lineages —Glaucophyta, Rhodophyta, Chloroplastida— were recovered with strong support by most subsets, as were other non-controversial clades such as SAR. However, Archaeplastida monophyly was not recovered by any of the 32 subsets of shared genes or taxa (except one with UFboot = 51%; Supplementary Table 1). Tree topologies obtained from the “gene overlap” datasets (with comparable genes but different taxa) clustered by source dataset that shared taxon sampling (Fig. 2a), suggesting that taxon sampling is a key factor in determining the overall tree topology. However, no taxa combinations that favored the monophyly of Archaeplastida was identified. In the case of topologies obtained from the “taxon overlap” datasets (with comparable taxa but different genes), the clustering by source dataset that shared gene sampling was less clear: some subsets converged to relatively similar topologies (e.g., BUR-BAU and BAU-BUR) whereas others did not (Fig. 2b). Trees containing only the 20 taxa shared by all four datasets had relatively similar topologies but differed in key relationships (i.e., deep relationships within Diaphoretickes). In both “gene overlap” and “taxon overlap” datasets, the trees obtained from the KAT dataset were the most scattered in the tree space (Fig. 2).

**Figure 1.**
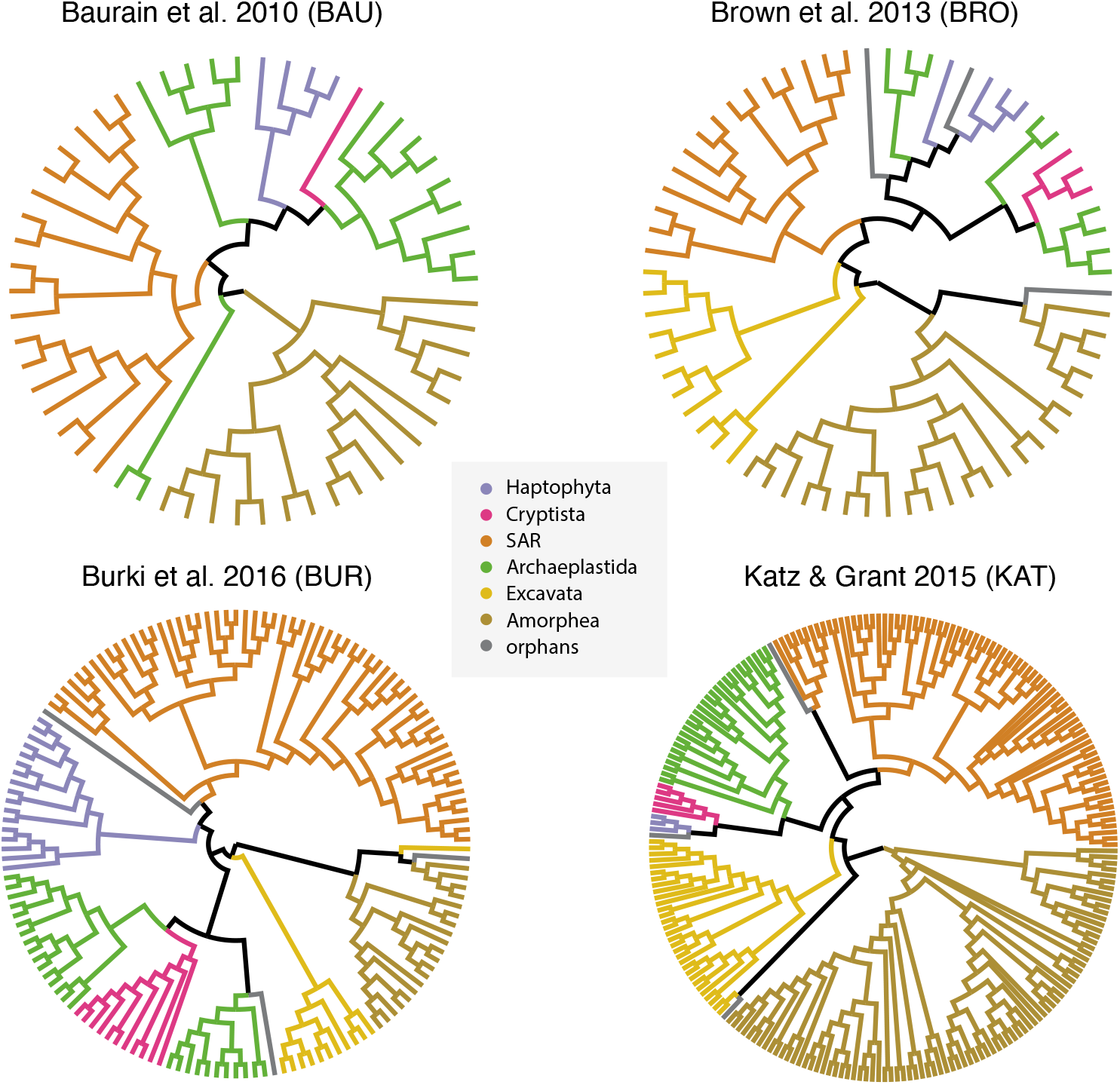
Maximum likelihood phylogenies inferred from the four original datasets (BAU, BRO, BUR, KAT) under empirical mixture models (IQ-TREE). Major eukaryotic lineages are shown (orphans: Telonemia, *Collodyction*, Picozoa).

**Figure 2.**
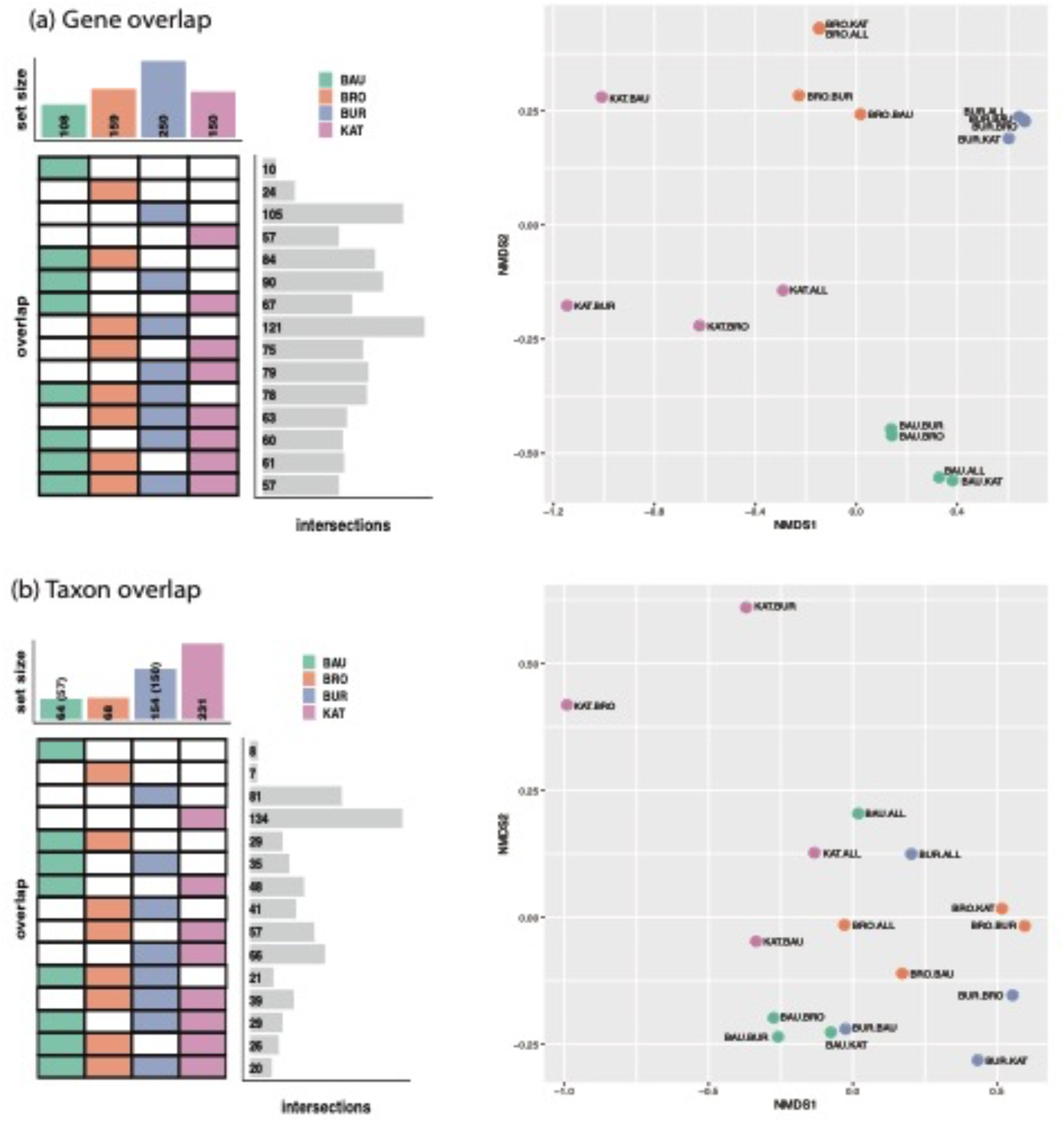
Taxon and gene overlap among the four analyzed datasets. Left graphs display the intersections of shared (a) genes and (b) taxa, whereas the right graphs show the topological distances among maximum likelihood trees from the corresponding overlapping datasets. Topological distances are measured as normalized Robinson-Foulds (nRF) distances between all pairs of trees and plotted after Kruskal’s non-metric multidimensional scaling (axes represent the inferred coordinates). Datasets are colored according to the source dataset.

### Undetected Paralogy Biases Deep Eukaryotic Relationships

Gene-wise likelihood scores (ΔGLS) showed widespread conflict in all four datasets regarding the relationships of PPE: about two thirds of the genes in BRO (93 vs. 66) and BUR (143 vs. 107) favored non-monophyletic PPE (χ^2^-test’s p < 0.05) whereas more genes supported the monophyly in BAU (48 vs. 60) and KAT (75 vs. 76) (Supplementary Table 2). For most genes, ΔGLS were small (< 10 for an average gene likelihood score of -lnL=13,481) but several outliers stood out in all datasets (Fig. 3). We investigated the 5% most deviant outliers in each dataset to identify causes for the conflicting signal regarding Archaeplastida monophyly. The majority of outliers from BAU, BRO, and BUR recovered PPE lineages clearly apart but with low support (e.g., *COPG2, DRG2, POLR3B* or *SARS* in BUR). Two loci (*RPL19* from BRO and *PSMC1* from BUR) recovered monophyletic Archaeplastida with low support. None of the inspected outliers in these three datasets showed obvious problems of paralogy or contamination, with two exceptions.

In BUR, the *UBA3* gene contained two divergent sequences that were likely contaminants or paralogs (*Plasmodium falciparum* and *Cyanidioschyzon merolae*; Supplementary Fig. 2). The removal of these two sequences did not significantly change the gene tree but switched the ΔGLS in favor of Archaeplastida monophyly. In BAU, the *EF2* gene displayed a strong signal against Archaeplastida monophyly (Fig. 3; Supplementary Fig. 3). Although we did not observe obvious contamination or paralogy problems, Glaucophyta was placed within Amorphea with strong support and away from a Rhodophyta + Chloroplastida clade, as reported before (Kim and Graham 2008). *EF2* alignment positions supporting the non-monophyly of PPE were not clustered, which could be expected from fused chimeric sequences (Supplementary Fig. 4). The removal of *EF2* from BAU did not alter the non-monophyly of Archaeplastida, but impacted other deep eukaryotic relationships. In particular, Glaucophyta was placed as sister to Chloroplastida + Haptophyta (Supplementary Fig. 5) and not as sister to all Diaphoretickes as in the original dataset (Fig. 1), consistent with the close affinity of Glaucophyta and Amorphea recovered by *EF2*. The *EF2* gene was also present in BRO and KAT, and while the gene trees consistently showed non-monophyletic PPE, ΔGLS were smaller (respectively, 3.06 and 0.68 vs. 50.10 in BAU). These differences could reflect a benefit of increased taxon sampling, but other factors such as rate differences decreasing the support for alternative topologies cannot be excluded (Walker et al. 2020).

In KAT, six out of the eight most extreme outliers strongly supported Archaeplastida monophyly (Fig. 3). A closer look revealed that all six genes had clear paralogy issues, often with a mix of nuclear, mitochondrial, and plastidial paralogs. Paralogs were confirmed by ML gene trees after the addition of nuclear, mitochondrial, and plastidial homologs from reference genomes (Supplementary Figs. 6-11). The removal of these paralogs from outlier loci in KAT was enough to break the monophyly of Archaeplastida recovered initially (IQ-TREE LG+C60+F+Γ4; Supplementary File 12).

### Larger Combined Datasets Have Higher Resolving Power

To overcome the limited phylogenetic signal in the four individual datasets, we assembled a larger combined dataset by accounting for shared genes and taxa, culminating in 311 genes and 344 taxa. This combined dataset was subjected to careful data curation to avoid the adverse effect of paralogy and contamination. We evaluated the gene support for Archaeplastida through ΔGLS, which again showed widespread conflict for Archaeplastida monophyly: 129 genes supported Archaeplastida monophyly whereas 182 genes did not (Supplementary Fig. 13). However, a close look at the 5% most extreme outliers revealed no obvious paralogy or contamination issues. To analyze this dataset, we derived three concatenated alignments after applying different site trimming strategies (COMB-UNTRIM, COMB-DIVPART, COMB-BMGE), and for each alignments we subsampled to 98 representative taxa for computational tractability of mixture models (see below).

**Figure 3.**
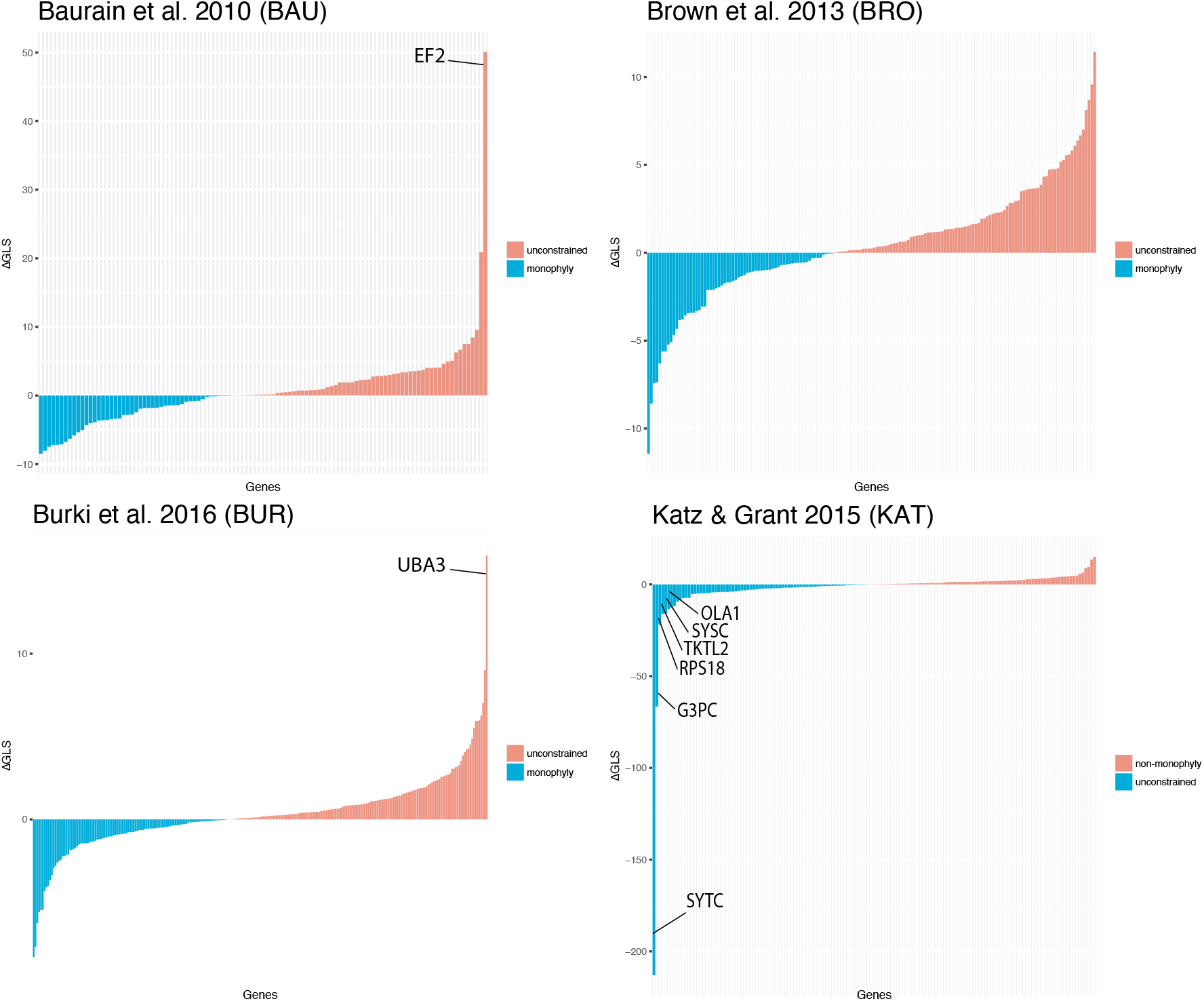
Outlier detection. Gene-wise log-likelihood score differences (ΔGLS) in support (blue) or against (red) the monophyly of Archaeplastida. Calculations follow Shen et al. (2017). Outliers discussed in the text are highlighted.

The completeness of all datasets were assessed with recently proposed alignment descriptive metrics (Wong et al. 2020) (Supplementary Table 3 and Supplementary Figs. 14 and 15). The individual datasets varied in their overall completeness (Ca) from 0.59 in KAT to 0.84 in BAU. The per-column completeness (Cc) was generally high, as expected from trimmed alignments. The per-row completeness (Cr) (i.e., taxa) was high for BAU and BUR but variable for BRO and especially for KAT. This was reflected in the higher proportion of shared amino acids between pairs of sequences (i.e., completely specified shared homologous sites or Cij) in BAU and BUR compared to BRO and KAT. In comparison, the combined datasets, especially those with the full taxon sampling, had generally lower overall completeness (Ca = 0.34–0.64) and the more aggressively trimmed datasets (BMGE) had higher matrix occupancy.

We also assessed the phylogenetic informativeness of all datasets by measuring the internal consistency among gene trees. Internal consistency was measured by normalized Robinson-Foulds distances (nRF) between gene trees and concatenated trees. Loci with limited signal are expected to produce gene trees that differ the most from the concatenated tree (stochastic error). Similarly, the presence of contaminants, paralogs, or very heterogeneous evolutionary patterns will also result in larger nRF. The combined datasets showed higher consistency (lower nRF) than any of the four original datasets, and were larger in terms of total alignment length and number of taxa. This pattern was maintained after correcting nRF for gene length, as shorter genes could be more prone to stochastic error (Supplementary Fig. 16). BAU, BRO, and KAT showed higher nRF than BUR, which was closer to COMB-BMGE, but the difference with KAT and BRO became smaller after correcting for gene length. Among the combined datasets, untrimmed alignments (COMB-UNTRIM) showed the highest consistency (lowest nRF), followed by COMB-DIVPART and COMB-BMGE. Internode certainty measures on the COMB-UNTRIM dataset identified substantial gene tree conflicts, particularly for deep branches (Supplementary Fig. 17). These conflicts did not derive primarily from short branches (Supplementary Fig. 18) as might be expected under high prevalence of incomplete lineage sorting.

### Aggressive Alignment Filtering Reduces Phylogenetic Signal

The effect of alignment trimming algorithms on recovering deep eukaryotic relationships was compared in detail by looking at the congruence between gene and concatenated trees, as well as the proportion of congruent and incongruent bipartitions in the combined datasets. While the distribution of nRF distances overlapped to a high degree, the mean nRF of COMB-UNTRIM was smallest (i.e., highest congruence), followed by COMB-DIVPART and COMB-BMGE (Supplementary Fig. 19a). COMB-UNTRIM also recovered the highest proportion of highly-supported congruent bipartitions whereas COMB-BMGE recovered more incongruent branches, but generally with low support (Supplementary Fig. 19b). COMB-DIVPART was indistinguishable from COMB-UNTRIM for short gene alignments (<400 amino acids) (Supplementary Fig. 19c). Despite the overall better performance of COMB-UNTRIM, COMB-BMGE and COMB-DIVPART performed best for a few genes, which were shorter than the average (Supplementary Fig. 19d-e).

### Monophyly of Archaeplastida and the Tree of Eukaryotes

The three combined datasets (COMB-UNTRIM, COMB-DIVPART, COMB-BMGE) were analyzed by ML under best-fitting site-homogeneous (LG(+F)+I+Γ4) and mixture models (LG+C60+Γ4), the latter with a reduced 98-taxon sampling for computational tractability. Under the site-homogeneous models, COMB-BMGE (Supplementary Fig. 20) recovered monophyletic Amorphea, including Opisthokonta, Amoebozoa, Apusomonadida, and Breviatea. As rooted with Amorphea, *Malawimonas* + *Collodyction* and Excavata were branching successively as sister to all other eukaryotes. Haptophyta was the sister group of a Telonemia + SAR clade (TSAR; Strassert et al. 2019). Archaeplastida was not recovered as monophyletic, with Cryptista being sister to Glaucophyta + Chloroplastida (with low support) to the exclusion of a Rhodophyta + Picozoa clade. COMB-DIVPART and COMB-UNTRIM (Supplementary Figs. 21 and 22) differed from COMB-BMGE in that fast-evolving Entamoeba (Amoebozoa) were recovered within long-branched metamonads (Excavata), likely due to long-branch attraction. COMB-UNTRIM further differed in the position of Telonemia, which was not recovered as sister to SAR but to Picozoa. This might be an artifact due to the relatively high proportion of missing data in both taxa (97,107 and 15,833 out of 202,042 aligned amino acid positions, respectively for Telonemia and Picozoa).

When analyzed under the better-fitting LG+C60+Γ4 mixture model, all 98-taxon combined datasets (COMB-UNTRIM-98, COMB-DIVPART-98, COMB-BMGE-98) converged to a very similar topology (Fig. 4). Compared with the 344-taxa datasets, Excavata were recovered in three successive lineages: (i) *Malawimonas*, (ii) Metamonada, and (iii) Discoba. Amoebozoa was recovered as monophyletic, indicating that the long-branch attraction artefact observed with site-homogeneous models was mitigated. Haptophyta was sister to SAR. Importantly, all 98-taxa datasets recovered the monophyly of Archaeplastida, with Cryptophyta as its sister group (Fig. 5 and Supplementary Figs. 23-25). The support for Archaeplastida increased with the total length of the alignments (COMB-BMGE-98: 46/88 UFBOOT/aLRT; COMB-DIVPART-98: 83/92; COMB-UNTRIM: 87/98). This was also the case for other relationships between major groups such as Haptophyta + SAR (Fig. 5 and Supplementary Figs. 23-25). Within Archaeplastida, Rhodophyta was sister to Glaucophyta + Chloroplastida with high support, which is consistent with the groupings observed in the analyses of the 344-taxa datasets (UFBoot > 98; SH-aLRT ≥ 96). The Bayesian analysis of COMB-BMGE-98 with the CAT-GTR model recovered identical deep eukaryotic relationships with strong support (BPP = 1.0), including monophyletic Archaeplastida sister to Cryptista, and Rhodophyta sister to Glaucophyta + Chloroplastida (Fig. 4 and Supplementary Fig. 26). CAT-GTR provided a better fit than LG+C60 (10-fold cross-validation score 1197.12 ± 81.5634), but as frequently seen in deep eukaryotic phylogenomic analyses (Burki et al. 2016; Brown et al. 2018; Gawryluk et al. 2019), the three MCMC chains failed to converge after >7,000 cycles (maxdiff = 1, meandiff = 0.0158). Notably, the lack of convergence was due to unresolved positions of long-branched amoebozoans and excavates, reflecting the difficulty of correctly placing these lineages, but all three MCMC chains agreed in the remaining bipartitions including Archaeplastida monophyly.

**Figure 4.**
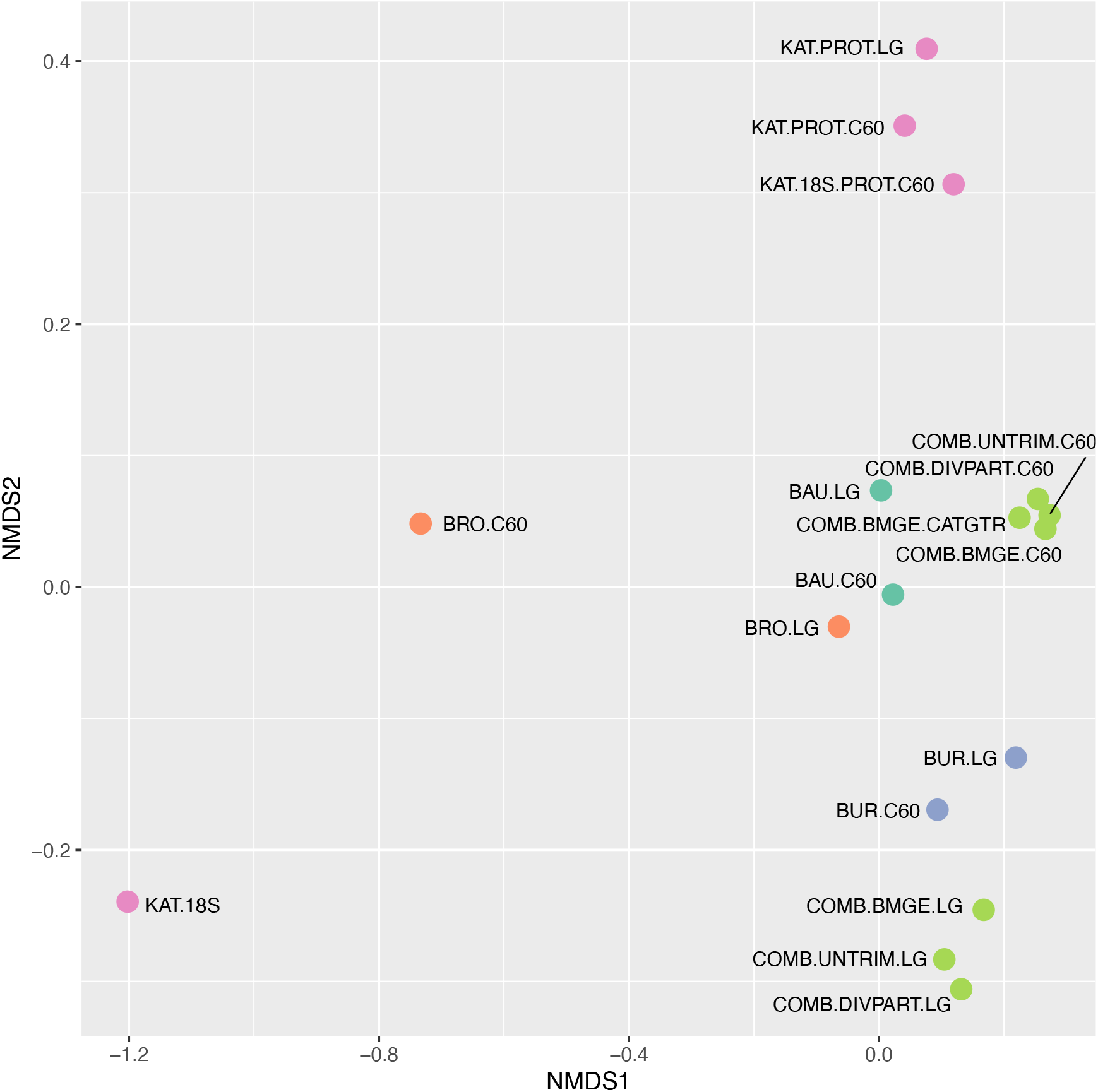
Topological distance among concatenated maximum likelihood trees from the original (BAU, BRO, BUR, KAT) and combined (COMB) datasets. Phylogenetic trees have been analyzed with both best-fit site-homogeneous (LG) and mixture (C60) models. KAT was analyzed in full and as separate 18S rRNA and protein partitions. Topological distances are measured as pairwise normalized Robinson-Foulds distances among shared taxa (nRF) and then plotted after Kruskal’s non-metric multidimensional scaling (axes represent the inferred coordinates).

**Figure 5.**
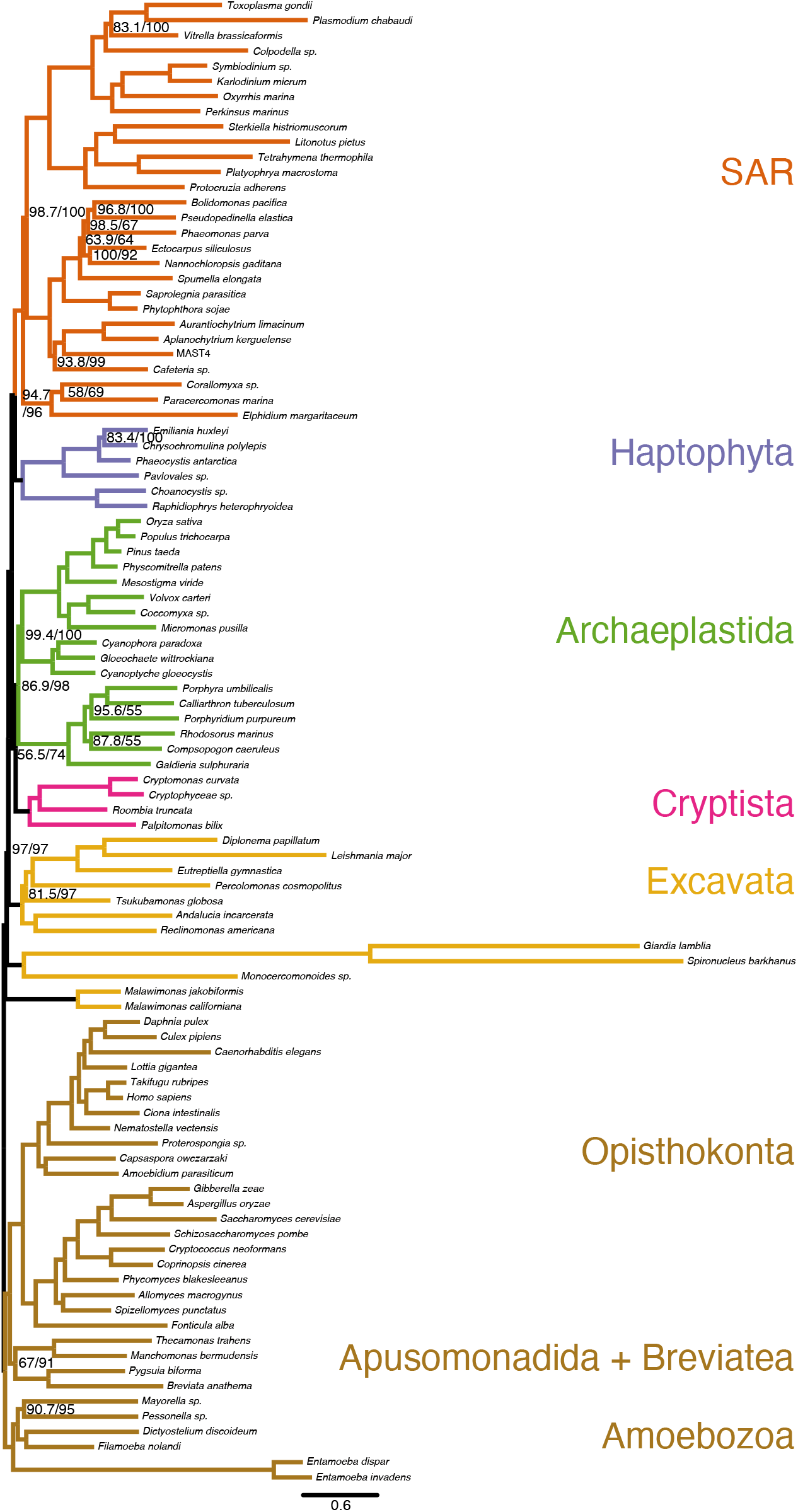
Maximum likelihood analysis of the combined (COMB-UNTRIM-98) dataset analyzed under LG+C60+I+Γ4 (IQ-TREE). Branch support is UFBoot/SH-aLRT; nodes without values received full support (100/100). Branch lengths are in expected substitutions per site.

## Discussion

### Resolving Recalcitrant Relationships in the Deep Phylogeny of Eukaryotes

Resolving the backbone of the eukaryotic Tree of Life is a complex endeavor requiring the inference of events dating back >1 Ga. Thanks to decades of phylogenetic research and more recently the advent of phylogenomics, the shape of the tree has been progressively refined, while some relationships have remained contentious (Burki et al. 2020). One such contentious grouping is Archaeplastida and its position within Diaphoretickes. Here, we have investigated the ancient origin of Archaeplastida and showed that recovering enough phylogenetic signal to resolve the monophyly of this group requires the assembly of larger datasets (in particular more genes), careful data curation for the removal of paralogs and contaminants, and the use of mixture models that better account for heterogeneous evolutionary patterns.

None of the four original datasets under study contained enough signal to resolve the phylogenetic affinities of PPE with confidence, nor to identify strongly supported alternatives. The identification of subsets of shared taxa and genes across the four datasets did not led to specific combinations of taxa or genes that helped recovering the monophyly of Archaeplastida. Heavily reducing taxon sampling to the 20 taxa shared by all four datasets produced more similar trees, but these were still inconclusive regarding PPE relationships. Heavily reducing gene sampling to the 57 genes common to all four datasets did not result in more similar trees, suggesting increased stochastic error with fewer loci. The poor signal for PPE relationships remained true after the identification of several paralogs that compromised phylogenetic accuracy (Fig. 3). In particular, the KAT dataset contained plastid paralogs (Supplementary Figs. 6-11) that artificially inflated the support for Archaeplastida due to the strong signal for monophyly in plastid genes (Baurain et al. 2010; Criscuolo and Gribaldo 2011; Ponce-Toledo et al. 2017; Sánchez-Baracaldo et al. 2017). The removal of the identified paralogs from this dataset led to the loss of the Archaeplastida monophyly recovered by the original dataset (Supplementary Fig. 12). In BAU, we found that the removal of one outlier locus (*EF2*) changed deep eukaryotic relationships. Although we could not identify obvious problems of paralogy or contamination, we observed a strong affinity between Glaucophyta and Amorphea, in agreement with previous studies suggesting hidden paralogy or horizontal gene transfer (Reeb et al. 2009; Atkinson and Baldauf 2010). These results confirm previous reports of phylogenomic datasets being driven by a handful of loci (Brown and Thomson 2017; Shen et al. 2017; Siu-Ting et al. 2019). Therefore, carefully dissecting the signal in phylogenomic datasets is crucial to identify problematic loci and biases and the distribution of phylogenetic signal. The development of new approaches to outlier identification (Walker et al. 2020) and automated pipelines for gene alignment curation (e.g., PhyloToL [Cerón-Romero et al. 2019]; PhyloFisher: https://pypi.org/project/phylofisher/) represent important advances for the assembly of large and accurate eukaryotic phylogenomic datasets.

Given the limited signal of the original datasets, we tested whether the combination of data improved the placement of PPE lineages. The combined datasets showed higher internal consistency than for any of the four source datasets (Supplementary Fig. 16), which we interpret as evidence for stronger genuine phylogenetic signal. Importantly, the strength of this signal was not associated with higher alignment completeness metrics (*sensu* Wong et al. 2020). The combined datasets also showed substantial conflict among genes with regards to the support of Archaeplastida (Supplementary Fig. 13) and deep eukaryotic relationships as a whole (Supplementary Fig. 17), although we did not identify obvious issues of paralogy or contamination in the outlier loci. Phylogenetic conflict measured as internode certainty was not preferentially associated with short branches, as expected under predominance of incomplete lineage sorting (Marcussen et al. 2014; Supplementary Fig. 18), suggesting that other sources of conflict, such as stochastic error or heterogeneous evolutionary patterns, are likely more prevalent (Bryant and Hahn 2020).

The phylogenetic analysis of the combined datasets under better-fitting mixture models converged to an overall consistent topology, in particular for the monophyly of Archaeplastida (Fig. 4). Mixture models can better account for heterogeneous evolutionary patterns in the data thereby reducing the risks of systematic errors (Rodríguez-Ezpeleta et al. 2007b; Philippe and Roure 2011; Wang et al. 2018). Simpler site-homogeneous models (LG) never recovered Archaeplastida, except in the presence of plastid paralogs in the KAT dataset. Model cross-validation showed that infinite profile mixtures (CAT), which infer amino acid profiles from the data, fit our combined datasets better than empirical mixtures with pre-defined number of profiles and weights (C60). Yet, both CAT-GTR (BI) and LG-C60 (ML) analyses reconstructed congruent tree topologies both consistently supporting the monophyly of Archaeplastida. The current implementation of CAT-GTR in PhyloBayes proved computationally challenging and failed to converge, as often seen in other studies of deep eukaryotic relationships (Kang et al. 2017 0515; Burki et al. 2016; Gawryluk et al. 2019). In this respect, the development of more efficient implementations of existing models (Dang and Kishino 2019) and new mixture models (Schrempf et al. 2020) should contribute to further resolving the eukaryotic tree of life.

### Effect of Alignment Filtering Algorithms

To understand the effect of alignment filtering in resolving the eukaryotic tree, we performed an empirical comparison between: (i) untrimmed data, (ii) the probabilistic algorithm Divvier (Ali et al. 2019), and (iii) BMGE, a commonly used block trimming method. In agreement with Tan et al. (2015), we found that untrimmed gene alignments retained more phylogenetic signal (lower nRF to the concatenated tree) than block-trimmed alignments (Supplementary Fig. 19a). Divvier, which was not assessed in Tan et al. (2015), was less accurate than no trimming but retained more signal than block trimming. Divvier’s reduced accuracy affected mostly longer genes (>400 aligned amino acids; Supplementary Fig. 19c). We note that aggressive block trimming does not only decrease the accuracy of gene trees, but can also affect the tree topology and support values of hundreds of concatenated loci (Supplementary Figs. 20–24). This is particularly the case when the phylogenetic signal is weak or confounded by substantial conflict. In the case of Archaeplastida, more aggressive trimming decreased the statistical support for its monophyly.

### Implications for Eukaryotic Evolution

Inferring the monophyly of Archaeplastida was only possible using the combined evidence from the four datasets, careful data curation, and the application of complex mixture models. Our analyses consistently recovered the monophyly of Archaeplastida with both ML and BI methods, in contrast to recent reports of conflicting ML and BI hypotheses (Brown et al. 2018; Gawryluk et al. 2019; Strassert et al. 2019). The monophyly of Archaeplastida has also been recovered in two recent studies (Lax et al. 2018; Price et al. 2019) although these are a minority among the datasets with a broad sampling of the eukaryotic diversity (Hampl et al. 2009; Baurain et al. 2010; Parfrey et al. 2010; Burki et al. 2012, 2016; Brown et al. 2013; Yabuki et al. 2014; Janouškovec et al. 2017). In our analyses, Cryptista was the sister group to Archaeplastida, a result that contrasts with previous studies that found it branching within Archaeplastida (Burki et al. 2016) or as sister to Haptophyta (Brown et al. 2018).

Within Archaeplastida, Rhodophyta was the sister group to Glaucophyta + Chloroplastida, in agreement with some recent phylogenomic analyses (Brown et al. 2018; Lax et al. 2018; Gawryluk et al. 2019; Price et al. 2019) and other studies that despite not recovering Archaeplastida, inferred a Glaucophyta + Chloroplastida clade (Burki et al. 2012, 2016; Brown et al. 2018). This contrasts with the earlier divergence of Glaucophyta favored by many plastid phylogenies (Ponce-Toledo et al. 2017; Sánchez-Baracaldo et al. 2017; Reyes-Prieto et al. 2018), even though the Rhodophyta-first hypothesis has also been recovered by plastid genes (Criscuolo and Gribaldo 2011; Lang and Nedelcu 2012). An earlier divergence of Chloroplastida has also been proposed based on plastid genes transferred to the nucleus during endosymbiosis (Deschamps and Moreira 2009).

### Origin and Evolution of Primary Plastids

In addition to further resolving the tree of eukaryotes, the recovery of Archaeplastida with nuclear data fills an important gap in our understanding of plastid origins. It is commonly assumed that primary plastids originated once in an ancestor of Archaeplastida (Reyes-Prieto et al. 2007; Gould et al. 2008; Löffelhardt 2014). This hypothesis of a single endosymbiosis was initially based on a similar double envelop surrounding both primary plastids and cyanobacteria, and the homology of the transit machinery for plastid protein import in all PPE. It also more easily explained some similarities of plastid genomes, such as the presence of a conserved set of genes and microsyntenic regions (Stoebe and Kowallik 1999), and the unusual tRNA-Leu group I intron (Besendahl et al. 2000). Endosymbiotically-derived gene clusters (Ku et al. 2015) and mosaic metabolic pathways (Reyes-Prieto and Bhattacharya 2007) have been interpreted as additional evidences for a common origin. Yet, all these evidence for a common origin are based on plastid data, but the similarities in plastid genomics and cell biology could at face value also be explained by alternative scenarios (e.g., independent or serial endosymbioses) in which hosts are not necessarily monophyletic (Supplementary Fig. 1) (Mackiewicz and Gagat 2014). Although typically considered less parsimonious than a single endosymbiosis, the possibility of serial endosymbioses has recently gained popularity in the case of the evolution of secondarily-derived plastid of red algal origin (Bodył et al. 2009; Stiller 2014). Similarly, secondarily-derived plastids of green algal origin are known to have independent origins (Rogers et al. 2007; Takahashi et al. 2007).

Therefore, it is critical to demonstrate not only the common origin of plastids, but also that host lineages of Archaeplastida are monophyletic, in order to make a strong argument in favor of a single primary endosymbiosis (Mackiewicz and Gagat 2014). As mentioned above, the monophyly of Archaeplastida has thus far only been sporadically recovered based on nuclear (host) markers, and it was unclear whether Archaeplastida was in fact polyphyletic or if stochastic and/or systematic errors in phylogenomic datasets have prevented the consistent inference of this clade. We have clarified this question by providing a well resolved eukaryotic tree with Archaeplastida consistently monophyletic, as well as describing a set of conditions that previously prevented the recovery of this group.

## Conclusion

In this study, we have investigated in detail the phylogenetic signal in four available datasets for one of the most pressing questions in eukaryote evolution: the monophyly of Archaeplastida. Neither the re-analysis of these datasets taken individually with better-fitting mixture models, nor various combinations of genes and taxa, provided enough signal to clarify the deep eukaryote relationships. It took the combination of the four datasets, together with a rigorous data curation pipeline and the application of complex mixture models, to recover enough information at this phylogenetic level. These analyses provided consistent support for the monophyly of Archaeplastida based on host markers, thus reconciling the evolutionary histories of plastids and hosts. This topology is compatible with the hypothesis of a single endosymbiotic origin of plastids in the Archaeplastida ancestor, establishing a firmer ground to better understand the early evolution in this important group of eukaryotes.

## Acknowledgements

We are grateful to D. Baurain and M. Brown for providing access to original data, to A. Rokas and X-X. Shen for sharing code, and to S. Whelan for insightful discussions. This work was supported by a fellowship from Science for Life Laboratory to FB. II was in part supported by a Juan de la Cierva – Incorporación postdoctoral fellowship (IJCI-2016-29566) from the Spanish Ministry of Economy and Competitiveness (MINECO). Computations were performed on resources provided by the Swedish National Infrastructure for Computing (SNIC) at Uppsala Multidisciplinary Center for Advanced Computational Science (UPPMAX) under projects SNIC 2017/7-151, SNIC 2019/3-305, and SNIC 2020/15-38.

## Data availability

The combined datasets and all phylogenetic trees are available from the Dryad Digital Repository: http://dx.doi.org/10.5061/dryad.[NNNN]

## Contributions

II and FB conceived the study; II and JFHS assembled datasets; II performed data analysis and wrote first draft; all authors contributed to the final text.

## Notes

### Competing Interest Statement

The authors have declared no competing interest.

